# Evidence provided by high-impact cell culture studies does not support authors’ claims

**DOI:** 10.1101/2021.04.12.439525

**Authors:** Ali Burak Özkaya, Caner Geyik

## Abstract

**Background:** Reliability of preclinical research is of critical concern. Previous studies demonstrate the low reproducibility in research and recommend raising standards to improve reproducibility and robustness. One understudied aspect of this quality issue is the harmony between the hypotheses and the experimental design in published work.

**Methods and findings:** In this study we focused on highly cited cell culture studies and investigated whether the claims of the study are backed with sufficient experimental evidence or not. We created an open access database containing all 282 claims asserted by 103 different high-impact articles as well as the results of this study. Our findings revealed that only 64% of all claims were sufficiently supported by evidence and there were concerning misinterpretations such as considering the results of tetrazolium salt reduction assays as indicators of cell death or apoptosis.

**Conclusions:** Our analysis revealed an alarming discordance between the actual experimental findings and the way that the manuscript is written to discuss them in highly cited cell culture studies. In order to improve quality of pre-clinical research, we require a clear nomenclature by which different cell culture claims are distinctively categorized, materials and methods sections to be written more meticulously and cell culture techniques to be selected and utilized more carefully.

## INTRODUCTION

There is an alarming concern regarding reliability of the published research findings [1]. This is particularly evident in preclinical studies as the clinical translatability is minimal [2]. This low efficiency in research has been discussed extensively in recent years and the lack of reproducibility and overall quality are agreed upon as the main culprits of the problem [3]. Reproducibility in preclinical research is estimated to be between 10% to 25% [4,5] and the cost of irreproducible research is calculated to be at least 28 billion USD/year in USA alone [6]. There are many factors contributing to this crisis including; lack of robustness, biased design, use of inadequate models (cell line and/or animal), underpowered studies (insufficient sample size), lack of proper controls (positive, negative), poor use of statistics and the absence of replication/confirmation studies [7]. It is important to note that these design problems often expand to questionable research practices such as p-hacking and cherry picking.

Scientists agree that the standards for publishing preclinical research must be raised in a way to encourage robustness and rigor [5,8]. Therefore, many aspects of the preclinical study design have been tacked by various studies over the years. However, the question of whether we can trust results of published preclinical studies remain at large. One aspect that has not been investigated before is the compatibility of the way the manuscript was written with the actual experimental design. More specifically, the relationship between the claims of the studies and the evidence provided to support these claims. In this study, we focused on cell culture research and investigated if the evidence provided by high-impact studies sufficiently supports the claims authors asserted in their manuscript. One of the first things we have noticed during our investigation was the inconsistency in the nomenclature. Many claims such as cytotoxicity, viability, growth, and proliferation were used interchangeably by the authors. Moreover, there were several publications in which only one type of evidence (tetrazolium reduction assay results) was provided to assert various claims. When we searched the literature to refine the consensus nomenclature, much to our disappointment, we could not find any. Many of these terms are not considered uncommon, unfamiliar, or vague enough to be defined in high-impact reviews or guidelines, or to be included in the glossary sections of the molecular biology, biochemistry and even cell culture textbooks. Therefore, we decided to define these terms ourselves mostly based on different sections of “Guidance Document on Good In Vitro Method Practices” by OECD [9] which was the only document we find that might be considered as a consensus nomenclature source. We then carried on our analysis accordingly. As a result, this study contains a nomenclature recommendation as well as the analysis of high-impact cell culture studies.

## METHODS

The study consisted of three phases. In phase one, we selected high-impact cell culture studies. In phase two, we identified the claims asserted by the authors as well as the evidence provided by them to support these claims. In the final phase we analyzed sufficiency of the evidence for each of the claims.

### Article Selection

We searched Web of Science (WOS) database (Clarivate Analytics) for studies that contain at least one of these keywords: “cytotoxicity, viability, cell death, growth inhibition, proliferation, or anti-cancer”. We included original research articles using *in vitro* techniques. The search string below was used in advanced search feature of WOS:

> WOS core database (TS= (“cytotoxicity” OR “viability” OR “cell death” OR “growth inhibition” OR “growth inhibitory” OR “proliferation” OR “anti cancer”) AND TS=(“cell culture” OR “in vitro” OR “cell line”)) AND LANGUAGE:(English) AND DOCUMENT TYPES: (Article)

Studies published in 2017 and 2018 were retrieved in 10.03.2020. We exported the data as an excel file, sorted the list based on citations received and selected the most cited 121 publications (each receiving at least 65 citations) as high-impact studies for further analysis. Upon investigation of the articles, we excluded 18 studies that do not contain a claim, were not carried out in cell culture, or were not original works. After exclusion of these ineligible studies, 103 studies were left for claim analysis.

### Claim Selection and Definitions

We identified seven distinct claims including: anti-cell, apoptosis, cell death, cell growth, cytotoxicity, proliferation, and viability in the studies. Using any of these terms to present or discuss a finding was considered as a claim.

We outlined our definitions for these claims as follows by using OECD guide GIVIMP [9]:

*Proliferation rate* represents how fast a group of cells divide over time. If the measurement was an end-point measurement, then it must be able to differentiate the decrease in division capabilities of the cells from cell death to provide sufficient evidence for proliferation rate. The reason is simple: if the treatment of question induces cell death in the treatment group, there would be fewer living cells (compared to untreated control) without a decrease in proliferation rate. Methods specifically focusing on replication rate (such as nucleotide incorporation) or measuring the number of viable cells without a treatment over time as well as real-time observations and proliferation markers were considered as sufficient evidence.

*Viability* represents the number of living cells. It is the broadest term since there is no specification regarding the factor affecting the number (such as proliferation rate or cell death). Any method directly measuring the number of living cells and methods measuring metabolic activity were considered as sufficient evidence for viability.

Since *cell death*, by definition, requires cells to die, end point analysis measuring the abundance of living cells cannot provide sufficient evidence for this claim as the measurement does not differentiate the decrease in number due to dying cells from slowed-down proliferation rate. Any assay measuring death-related alterations such as membrane integrity or cell-death specific markers was considered to provide sufficient evidence.

*Apoptosis* is a form of programmed cell death which has well established and characterized distinct features. The methods that are capable of demonstrating the changes in these features such as phosphatidylserine exposure on the outer membrane, DNA fragmentation, morphological changes and molecular switch responsible in apoptosis were considered as sufficient evidence to show apoptosis.

*Cytotoxicity* indicates being toxic to cells. Being toxic itself is a broad term and there are conflicting definitions in use. We decided to consider it as cell death instead of decreased viability as the most widely accepted capability of a toxic agent is killing (as in cytotoxic T cells and cytotoxic chemotherapy), and accepted evidence indicating cell death as sufficient to prove cytotoxicity. This decision had an impact on the final analysis which might be considered controversial as many of the articles might have used the term to represent viability decrease. We addressed this in results section. We decided to consider *anti-cell* (which was asserted just once) as if it was referring to cell death as well.

*Cell growth* may indicate either proliferation rate or the size change of cells depending on the definition embraced. Since, there already is a term representing proliferation rate as the name implies, we first considered to accept it as a measure of increased cell size. However, after looking at the articles in our list, we realized that the term almost exclusively used to indicate proliferation rate and consequently we embraced that definition in our analysis.

### Database Construction

We constructed a database in Airtable to carry out evidence analysis. Information from WOS database such as “article name”, “DOI”, “citation count”, “journal name” as well as our parameters of interest such as “claim”, “evidence”, “method”, “sufficiency of evidence”, and “subject area” were entered for every article investigated.

Here, “method” represents scientific methods used in the study whereas “evidence” is defined as a supergroup of methods measuring same biological phenomenon. For example, two separate methods such as lactate dehydrogenase (LDH) activity assay and PI both of which measure membrane damage as an indicator of cell death were classified into “membrane integrity” evidence supergroup. Similarly, various tetrazolium and resazurin reduction assays were considered to provide “dehydrogenase activity” evidence which is an indicator of cellular metabolic activity.

We have also divided the studies in two notional groups of “subject area” first being “Biochemistry, Molecular Biology, Genetics, and Medicine” and second “Chemistry, Chemical/Biomedical Engineering, and Materials Science” based on field information provided by WOS.

Database is accessible via the link: https://airtable.com/shrClTE87e1l28ExG

### Evidence Analysis

Evidence was analyzed for each of the claims by a case-by-case approach. We refer to the definitions we have embraced and the OECD guide GIVIMP [9] to determine if the measured parameter provides sufficient evidence for that claim. Table 1 summarizes the detected claims and the evidence considered as sufficient or insufficient. It is important to note that if there were multiple evidence for a single claim, we focused on the strongest evidence as well as the strength of the combination of the evidence. Therefore, in the table if the evidence is listed under “insufficient” it means it does not provide sufficient evidence by itself. For example, even though expression of Blc-2 members (such as Bax/Bcl-2 ratio) is commonly investigated along with other markers of apoptosis, it is classified as insufficient because it does not provide proper evidence for apoptosis by itself.

**Table 1.** List of sufficient and insufficient evidence for claims.

| Claim | Sufficient | Insufficient |
| --- | --- | --- |
| Anti-cell |  | Dehydrogenase activity |
| Apoptosis | Caspase-3 activity, Cell morphology, DNA fragmentation, Phosphatidylserine exposure | Dehydrogenase activity, Membrane integrity, Bcl-2 Family expression |
| Cell Death | Membrane integrity, Phosphatidylserine exposure | Cell count, Dehydrogenase activity, Esterase activity |
| Cell Growth | Cell count, Colony formation, Nucleotide incorporation | Dehydrogenase activity |
| Cytotoxicity | Membrane integrity, Phosphatidylserine exposure, | Dehydrogenase activity |
| Proliferation | **Cell count\***, **Colony formation\***,<br>**Dehydrogenase activity\***, **DNA amount\***, Dye<br>inclusion, Ki67 expression, Nucleotide<br>incorporation, Phospho-histone H3 | Actin, **Cell count\***, **Colony**<br>**formation\***, **Dehydrogenase**<br>**activity\***, **DNA amount\***,<br>Esterase activity, Signaling<br>Pathways |
| Viability | ATP amount, **Cell count\***, Dehydrogenase<br>activity, Dye inclusion, Esterase activity,<br>**Membrane integrity\***, Total protein | **Cell count\***, **Membrane**<br>**integrity\***, Signaling Pathways,<br>Unknown |

\* The evidence is either sufficient or insufficient for given claim depending on the study design.

There were some cases where the same type of evidence may be sufficient or insufficient based on the experimental design. Viability indicators such as cell count, colony formation, DNA amount, and dehydrogenase activity were considered to provide sufficient evidence for proliferation only if there is no treatment (hence no reason for a change in cell death ratio) and the study is designed in a way to observe proliferation of these untreated cells. Moreover, if the method utilized to obtain the evidence was not mentioned in materials and methods section of the article, we then classified it as insufficient (independent from the actual evidence) which was the case for cell count providing insufficient evidence for viability and proliferation (article id#7 from Airtable database). Another such deviation was the membrane integrity as an evidence for viability. Cell death marker PI was used as an evidence for viability in one article (article id#66 from Airtable database) which was considered as insufficient. All other articles (6 articles in total) claimed viability by providing different membrane integrity assays (including 7-AAD, GhostDye, Trypan Blue and Zombie UV) all of which measure viable cells.

### The Effects of the Journal and Subject Area on Evidence Sufficiency

After the sufficiency of evidence was assessed for each claim, we then sought to find whether being published in a specific journal or in a subject area would affect sufficiency rate. Seven journals were selected for analysis as they meet our criteria of having at least 10 claims, namely *Biomaterials*, *Cell*, *Cell Death Dis*., *Mol*. *Cancer*, *Nature*, *Oncotarget*, *Ann*. *Biomed*. *Eng*. (97 claims out of 282, 34.40%). Every journal was compared to the complete data set excluding themselves. Two subject areas: “Biochemistry, Molecular Biology, Genetics, and Medicine” and “Chemistry, Chemical/Biomedical Engineering, and Materials Science” were compared to each other. Fisher’s exact test (2-tail) were used in all comparisons.

## RESULTS

We have investigated 282 claims asserted by 103 different high-impact articles. We identified 7 unique claims supported by 20 types of evidence all of which was obtained via 40 different methods. Details are presented in the database (https://airtable.com/shrClTE87e1l28ExG)

The most common claim was proliferation rate changes with 85 claims, followed by viability, apoptosis, cytotoxicity, cell death, growth changes and anti-cell activity (Figure 1A). Upon investigation, we considered the evidence of 102 claims (36%), which was asserted by 66 different studies (64%), as insufficient (Figure 1B). Claims of cytotoxicity (11 sufficiency rate in 37 claims, 30%), proliferation (35 in 85, 41%), and cell growth (10 in 18 claims, 56%) particularly lacked proper evidence. Viability (66 in 72, 92%), apoptosis (46 in 51, 90%), and cell death (14 in 18, 78%), on the other hand, were more frequently claimed with proper evidence (Figure 1A).

**Figure 1.**
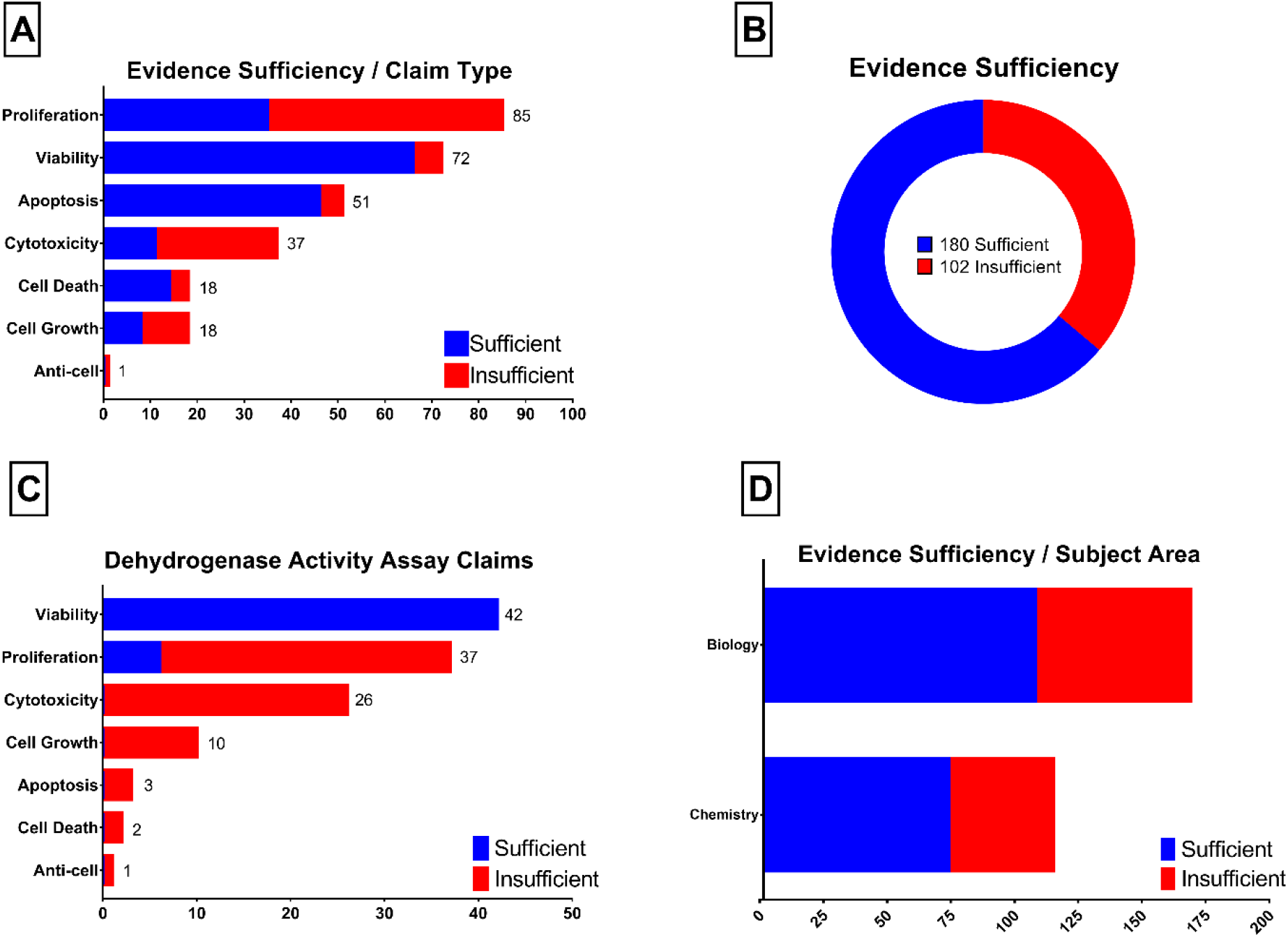
Evidence sufficiency analysis results. 1A. Evidence sufficiency according to the claim type; 1B. The ratio of claims that are supported by sufficient evidence or not; 1C. Evidence sufficiency of claims supported by dehydrogenase activity assay; 1D. Evidence sufficiency according to the article subject area

When evaluating claims, we had to make assumptions on behalf of the researchers by assigning specific definitions to the claims which is listed in the materials and methods section of this manuscript. Cytotoxicity was most likely to be used differently in articles we analyzed, possibly affecting the results of our analysis. If we had considered cytotoxicity as a measure of viability instead of cell death, all the cytotoxicity claims would have had sufficient evidence increasing the overall sufficiency rate from 64% to 73%.

Measurement of dehydrogenase activity was by far the most common type of evidence. 121 different claims (44% of all claims) including viability, proliferation, cytotoxicity, cell growth, cell death, anti-cell and even apoptosis put forward dehydrogenase activity findings as evidence (Figure 1D). The most common method used to measure dehydrogenase activity was tetrazolium reduction assay supporting 115 different claims (41% of all claims). Measurement of dehydrogenase activity was followed by measurement of phosphatidylserine exposure (31 claims) and membrane integrity (29 claims).

While the results of dehydrogenase activity assays were interpreted correctly only in 73 claims (73 in 121, 60%), the second most common evidence, measurement of phosphatidylserine exposure, was interpreted correctly (as an indicator of apoptosis) in all related claims (31 in 31, 100%). Similarly, measurement of membrane integrity was mostly correctly utilized as evidence for cell death claims (27 in 29, 93.1%) (Figure 1C)

We also analyzed whether the subject area or the journal that the article was published in might be an indicator of evidence claim relationship. According to our analysis, the subject area did not have a significant influence over evidence sufficiency (Figure 1D). Similarly, amongst the most frequently appeared journals in our database none of them demonstrated a significant difference when compared the overall claims supported by sufficient evidence.

## DISCUSSION

Our findings reveal a discordance between the claims and the evidence of high-impact cell culture studies. This is especially evident in studies utilizing the findings of tetrazolium reduction assay alone to support various claims. Striking examples include article id#9 claiming viability, proliferation, cytotoxicity and apoptosis changes, and article id#26 claiming viability, proliferation, growth changes and anti-cell activity with results from this assay. This is partly because these assay kits are advertised by their manufacturers as a tool to measure viability, cytotoxicity, proliferation and growth. Combining this with being relatively easier to perform and affordable leads to these assays being perceived as a one-size-fits-all solution by research groups wishing to avoid more complicated cell culture techniques. However, this reductionist approach makes it difficult for the findings obtained from the study to provide meaningful answers to the research questions of the article. Even though the reduction of cellular metabolic activity is not a clear indicator of apoptosis or cell death, since that statement exists in a high-impact article, there is a good chance that it will be cited as such as well.

In fact, the articles we analyzed were cited more than 9000 times as of March 2021 (within two to four years). Even though the claims without sufficient evidence may not be the reason for citation in most of the cases, the impact of such studies with unreliable findings on preclinical research is undeniably large. Many agree that we need strategies in place to improve the standards for pre-clinical research [5,8]. As a part of this process, we need clear and distinct definitions for terms corresponding to the claims asserted in the articles. In this work we offered a nomenclature recommendation by which the most common claims in cell culture studies may be distinctively expressed. Moreover, scientists both as authors and reviewers are required to ensure that the experimental designs reflect the research questions asked. As the findings of our study indicate, we believe a more meticulously written materials and methods section and careful selection and utilization of cell culture techniques are critical to raise robustness and overall quality of preclinical research.

## ACKNOWLEDGEMENTS

We would like to thank Yasemin SEVAL ÇELİK for her contributions in revising this manuscript.

